# pCASPR: A Single-Plasmid Platform for Scarless Bacterial Genome Editing

**DOI:** 10.1101/2025.08.05.668820

**Authors:** Pietro Ridone, Nathan Paul, Alexander R. Kelly, Dominic Glover, Thomas Huber, Matthew A. Baker

## Abstract

Scarless bacterial genome editing enables the construction of stable chromosomal genotypes free of residual selectable-markers or recombination site scars, but practical workflows can be limited by plasmid handling, guide retargeting, curing, and host-specific implementation. CRISPR-Cas9/lambda-Red systems such as no-SCAR, pREDCas9/pGRB, and pCas/pTargetF established efficient donor-templated editing in *Escherichia coli*, but many workflows still require multi-plasmid handling, repeated guide-plasmid construction, separate curing steps, or host-specific redesign. Here we describe pCASPR (CRISPR-Assisted Single-Plasmid Recombineering), a compact single-plasmid CRISPR-Cas9/lambda-Red platform designed to simplify retargeting, editing, and plasmid clearance while preserving the familiar counterselection-recombineering logic.

The system consolidates inducible Cas9/sgRNA counterselection, arabinose-inducible lambda-Red recombination, Golden Gate/GFP-dropout guide exchange, antibiotic selection, and temperature-sensitive curing in one vector. We devised this single-plasmid architecture from the widely used no-SCAR system to retain user compatibility and keep workflow integration seamless, while adding the benefits of a single-plasmid design and streamlined retargeting version for *E. coli*. In editing assays across 4 loci, pCASPR showed lower escape frequencies than the parental 2-plasmid system, with matched or improved editing efficiencies. pCASPR also enabled an *mVenus* knock-in at the *motAB* locus that was not recovered with no-SCAR. Guide exchange, edit recovery, counterselection outcome, and plasmid loss could be followed within the same workflow, with a saving of five days compared with the original no-SCAR protocol. The same single-plasmid design was implemented in *Shewanella oneidensis*, where the vector enabled the knock-in of a SpyTag fragment at the *mtrC* gene.

Together, these data define the design principles and demonstrate the utility of modular single-plasmid workflows for scarless bacterial genome editing, intended for easy adoption across bacterial species.

## Introduction

A clean chromosomal edit is often the decisive genetic test in bacteria. When a phenotype is assigned to a deletion, allele replacement, or short insertion, the chromosome should carry the intended change without leaving an antibiotic cassette, plasmid-borne copy, or residual recombination site. This is especially important for iterative strain construction and experimental evolution, quantitative genotype-phenotype comparisons, and loci where local sequence context affects expression or function.

Lambda-Red (λ-Red) recombineering made chromosomal editing in *Escherichia coli* rapid and accessible by using phage recombination functions to incorporate linear DNA donors through short homology arms {Murphy, 1998 #1; Zhang, 1998 #2}. The Datsenko-Wanner method built on this logic by pairing inducible lambda-Red expression with PCR-amplified resistance cassettes, allowing targeted gene disruption and later marker removal by Flp-FRT recombination {Datsenko, 2000 #3}. Cassette excision leaves flippase recognition target (FRT) scars and repeated edits require iterative marker management {Datsenko, 2000 #3}. This can complicate later rounds of engineering.

Scarless recombineering edits a defined chromosomal allele without a residual selectable-marker footprint. CRISPR-Cas9 counterselection changed the selection problem: instead of selecting the donor cassette, Cas9 can be directed to cut the unedited allele, enriching cells that have incorporated the intended donor sequence {Jiang, 2013 #4}. The no-SCAR system combined Cas9 counterselection with lambda-Red recombineering to recover scarless *E. coli* edits, and its protocol established a practical reference workflow {Reisch, 2015 #5; Reisch, 2017 #6}. Related CRISPR/lambda-Red approaches showed the same basic principle could support chromosomal gene replacement in *E. coli* {Pyne, 2015 #7}.

Several established *E. coli* CRISPR editing systems use two-plasmid architectures. The compact benchmark is pREDCas9/pGRB, where pREDCas9 carries Cas9, lambda-Red, and a plasmid-curing module, while pGRB supplies the retargetable guide RNA during editing {Li, 2015 #8}. Although powerful and compact, this remains operationally a two-plasmid workflow, because each new target requires reconstruction of the guide plasmid and subsequent removal of that plasmid after editing. The widely used pCas/pTargetF system similarly separates Cas9/lambda-Red functions from target-specific guide expression, enabling iterative multigene editing but requiring maintenance, retargeting, and curing of a second plasmid {Jiang, 2015 #9}. Later derivatives, including pEcCas/pEcgRNA, improved strain compatibility and guide-expression control but retained the same basic two-plasmid logic {Li, 2021 #10}. More recent *E. coli* editing platforms continue to refine guide control, survivor suppression, and iterative recovery, underscoring that practical usability depends not only on Cas9 activity but also on how retargeting, plasmid burden, and post-editing clearance are handled {Fang, 2024 #11; Lim, 2024 #12; Li, 2023 #13}.

Here we describe pCASPR (CRISPR-Assisted Single-Plasmid Recombineering), a platform whose defining feature is its single-plasmid architecture. pCASPR consolidates Cas9/sgRNA counterselection, lambda-Red recombineering, antibiotic selection, and temperature-sensitive replication for plasmid curing into one vector, so the one plasmid is transformed, retargeted, induced and removed. This eliminates the need for a separate guide plasmid and also for a second round of CRISPR-mediated curing, while maintaining the familiar counterselection-recombineering logic. In contrast, No-SCAR requires two plasmids to be introduced sequentially, as they were not able to be co-transformed, which was attributed to toxicity from leaky Cas9/sgRNA expression {Reisch, 2015 #5}. The pCASPR plasmid described here was derived from the two no-SCAR plasmids, pCas9cr4 and pKDsgRNA. Guide RNA exchange is facilitated by Golden Gate cloning combined with a visual GFP-dropout guide exchange, which also can be used to determine whether temperature-sensitive curing works in new host species.

To test whether this plasmid architecture is usable beyond *E. coli*, we adapted it for *Shewanella oneidensis* (pCASPR*-So*) by exchanging the antibiotic marker and origin of replication while retaining the same single-plasmid editing logic. Recombineering and CRISPR/Cas9 counterselection have already been established for *S. oneidensis*, and their performance varies based on transformation efficiency, recombinase choice, plasmid replication, selectable markers, and expression strength {Cao, 2019 #14; Corts, 2019 #15; Corts, 2019 #16; Suzuki, 2020 #17}. Our aim with *S. oneidensis* was to demonstrate the single-plasmid design could be transferred to a new species by swapping in host-compatible modules.

Specifically, we (i) construct pCASPR and validate its inducible Cas9/sgRNA counterselection and temperature-sensitive curing in *E. coli*; (ii) compare its editing performance with the parental two-plasmid no-SCAR system at multiple chromosomal loci; and (iii) adapt the architecture to *S. oneidensis*, demonstrating SpyTag knock-in at the native *mtrC* locus. By reducing editing to a single retargetable plasmid, this architecture is designed to shorten cloning and strain-construction workflows, simplify retargeting and the recovery of plasmid-free edited strains.

## METHODS

### Strains, plasmids, and growth conditions

*E. coli* RP437 was used for the pCASPR-*motAB* editing experiments, including *motAB* deletion and fluorescent-protein knock-in assays. *E. coli* BL21(DE3) was used for the pCASPR multi-locus counterselection and editing assays. *Shewanella oneidensis* MR-1 was used for pCASPR*-So-mtrC* counterselection and *mtrC*-SpyTag editing experiments using pCASPR*-So-mtrC-*HDR. Plasmids, N20 targeting sequences and donor DNA sequences are listed in Supplementary Table S1; oligonucleotide primers are listed in Supplementary Table S2.

Unless otherwise stated, *E. coli* cultures carrying pCASPR (Spec^+^) or no-SCAR plasmids (Spec^+^/Cm^+^) were maintained in LB medium at 30°C. Temperature-sensitive plasmid curing was performed at 42°C in non-selective LB. Antibiotics were used as appropriate for each plasmid: spectinomycin (Spec) at 50 μg/mL for pKDsgRNA and pCASPR-derived constructs, chloramphenicol (Cm) at 25 μg/mL for pCas9cr4 in the no-SCAR workflow, and kanamycin (Km) at 25 μg/mL for pCASPR-*So* constructs. Anhydrotetracycline (aTc) was used at 200 ng/mL unless otherwise stated, and L-arabinose was always used at 0.2% (w/v).

Soft-agar motility assays were performed on an LB medium substrate (Yeast Extract, Tryptone) adjusted for agar content (0.25% Bacto agar) and 85 mM NaCl. Individual colonies were transferred from selective plates onto the soft-agar swim plate using a pipette tip, then incubated at 30°C for 20 h. Positive motile clones were assessed by the presence of a halo surrounding the site of inoculation (swim rings) while non-motile clones grew as a solid colony at the point of inoculation.

### pCASPR plasmid construction and validation

The first-generation single-plasmid editor, pCASPR-*motA*, was assembled from the no-SCAR-derived plasmids pCas9cr4 and pKDsgRNA (Addgene #62655 and #62656; Fig. 1B, SI Fig. 2A, {Reisch, 2015 #5}. The pCas9cr4-derived fragment supplied the *tetR*/tetO-regulated Cas9 module, and the pKDsgRNA-derived fragment supplied the sgRNA expression module, *araC*/pBAD-regulated lambda-Red recombination genes, spectinomycin resistance, and a temperature-sensitive pSC101-derived replication module. PCR fragments were generated with primers carrying 20 bp overlap regions and assembled by overlap-based cloning using Takara In-Fusion Snap Assembly (Takara Bio). Assembled products were transformed into *E. coli* and candidate plasmids were screened by diagnostic restriction digestion and sequencing. The pCas9cr4-derived fragment was approximately 5 kb, the pKDsgRNA-derived fragment was approximately 7 kb, and the completed pCASPR plasmid was linearized with NotI to yield an approximately 12 kb band (SI Fig. 2). The assembled junction and guide region were verified by Sanger sequencing, and final plasmid sequences were verified by whole-plasmid sequencing (SI Fig. 2B). The full pCASPR-*motA* sequence file is available in SI.

**Figure 1.**
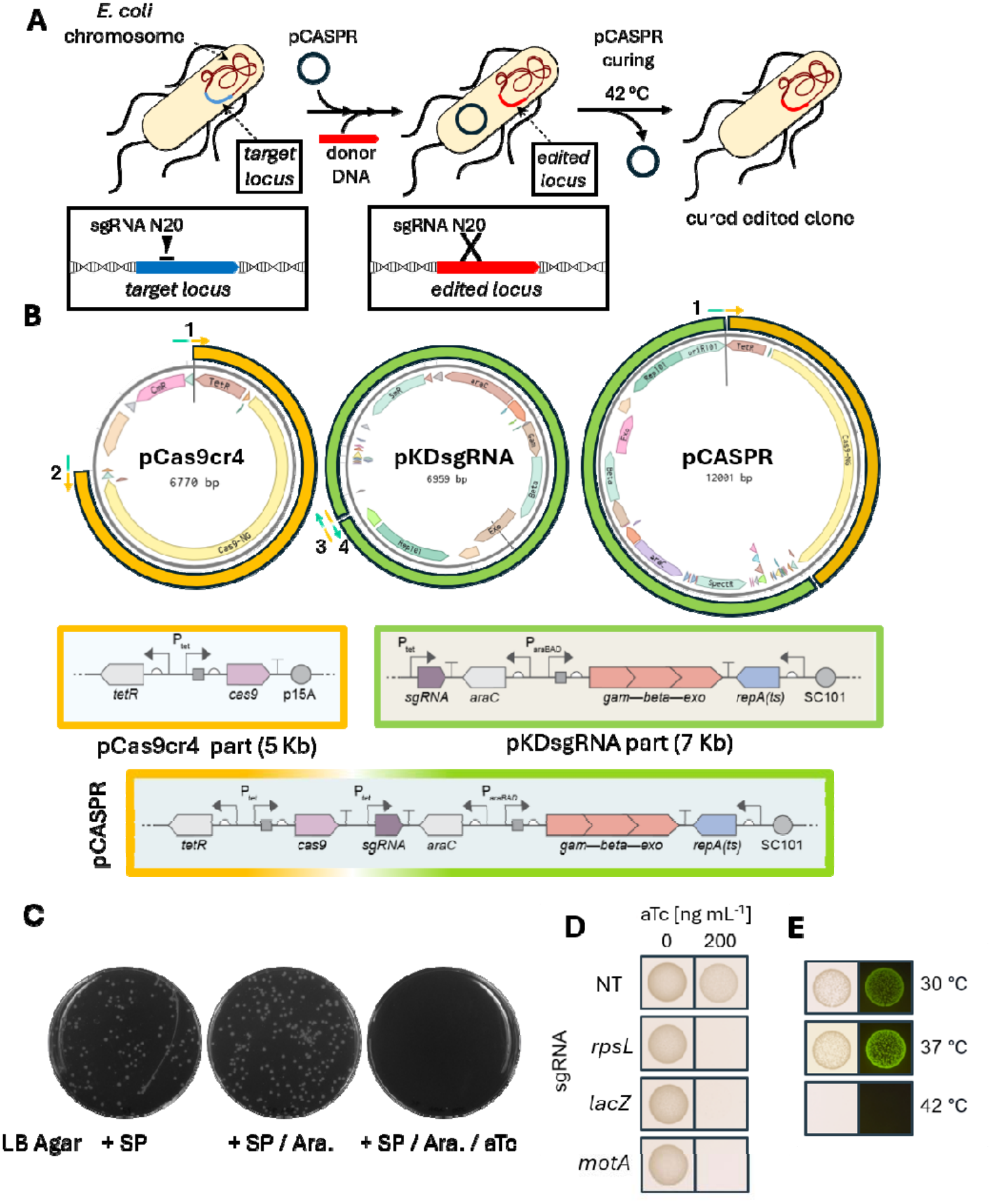
Construction and validation of pCASPR, a single-plasmid CRISPR/Cas9-lambda-Red genome-editing vector. (A) Schematic workflow for genome editing in *E. coli* using pCASPR. The pCASPR plasmid and double stranded donor DNA template for homologous recombination (dDNA) are sequentially introduced into the target strain to induce a dsDNA break near the sgRNA N20 hybridization site and homology directed repair (HDR) using the supplied dDNA. Successful integration of dDNA during HDR removes the N20 recognition site and prevents further dsDNA breaks. Failed integration results in cell death (counterselection). After screening for positive edits, the pCASPR plasmid can be removed from the selected clone by growth at 42°C (curing). (B) Assembly strategy for pCASPR. The no-SCAR-derived pCas9cr4 and pKDsgRNA plasmids {Reisch, 2015 #5} were used as sources of the Cas9 counterselection and lambda-Red recombineering modules, respectively. The pCas9cr4-derived region supplies the *tetR*/tetO-regulated *cas9* cassette, whereas the pKDsgRNA-derived region supplies the sgRNA expression module, *araC/*pBAD-regulated lambda-Red genes (*gam, bet, exo*), spectinomycin resistance, and temperature-sensitive replication/curing features. (C) Functional test of plasmid maintenance and induction. Cells carrying pCASPR-*motA* were plated on LB plus spectinomycin, LB plus spectinomycin and arabinose, and LB plus spectinomycin, arabinose, and aTc. Growth under spectinomycin selection shows plasmid maintenance, while loss of recovery after combined arabinose and aTc induction is consistent with donor-free Cas9/sgRNA counterselection at the chromosomal *motA* target. (D) aTc-inducible Cas9/sgRNA activity tested with a non-targeting guide and guides directed against *rpsL, lacZ*, or *motA*. The non-targeting guide retained growth after aTc induction, whereas target-matched guides reduced colony recovery, consistent with guide-dependent cleavage of chromosomal targets. (E) Temperature-sensitive plasmid maintenance/curing assay using a pCASPR-GFP:DO. Growth and GFP signal were retained at permissive temperatures, whereas growth/fluorescence was lost at 42°C under plasmid-selective conditions, consistent with the temperature-sensitive origin used for plasmid clearance.

**Figure 2.**
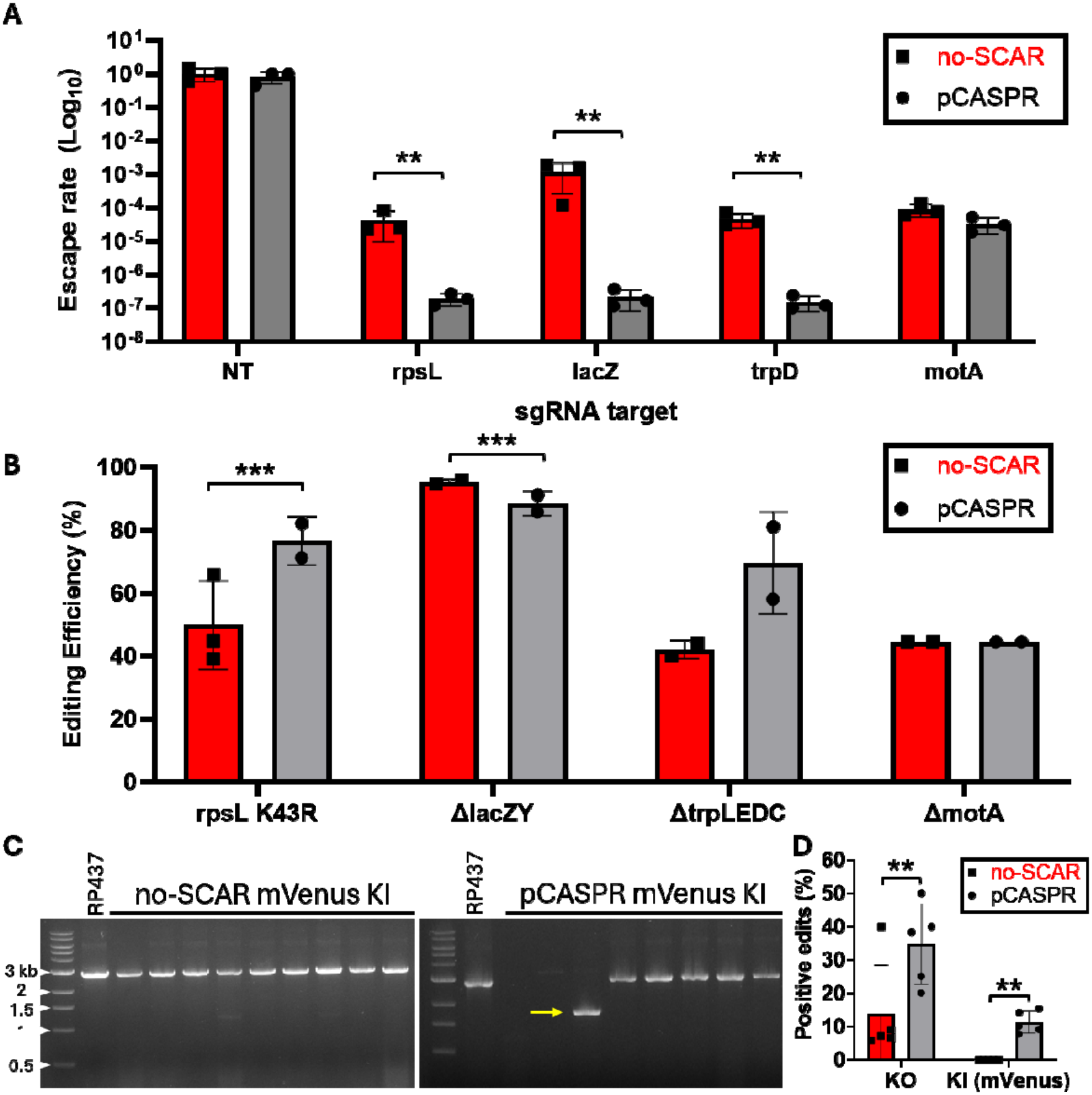
pCASPR supports retargetable multi-locus editing in *E. coli*. **(A)** Counterselection survival for no-SCAR and pCASPR after aTc induction of Cas9/sgRNA in *Ec* BL21. A non-targeting guide was used as the permissive control, and target-specific guides were directed against *rpsL, lacZ, trpD*, or *motA*. Colony recovery was high for the non-targeting guide and strongly reduced for chromosomal target-matched guides. Escape frequencies were calculated as the ratio of CFU/mL recovered on selective (+aTc) and non-selective (−aTc) plates. Bars represent mean ± SD from three independent biological replicates per treatment (N = 3). Statistical comparisons between pCASPR and No-SCAR were performed using unpaired Welch’s t-tests on log□□-transformed escape frequencies, with Holm correction for multiple comparisons. Adjusted p-values were: NT, p = 0.7351 (no significant difference); rpsL, ** p = 0.00461; lacZ, ** p = 0.00215; trpD, * p = 0.00144; and motA, p = 0.1180 (no significant difference). **(B)** Editing efficiency among screened colonies for no-SCAR and pCASPR at four chromosomal targets using donor-templated editing: *rpsL* K43R, Δ*lacZY*, Δ*trpLEDC*, and Δ*motA*. Editing efficiency was calculated as the percentage of screened colonies carrying the intended genotype, as determined by screening 18 random colonies from each of 2 independent editing experiments. pCASPR recovered edited colonies at all tested loci, with locus-dependent performance relative to no-SCAR. Data are plotted as mean ± SD; red, no-SCAR; grey, pCASPR. Editing efficiencies were determined from rpsL K43R (No-SCAR: N = 3, 236 StrepR/462 Spec; pCASPR: N = 2, 236 StrepR/303 Spec; \*\*\**p* < 0.001), lacZ (No-SCAR: N = 2, 206 Blue/4,690 White; pCASPR: N = 2, 113 Blue/958 White; \*\*\**p* < 0.001), trpLEDC (No-SCAR: N = 2, 45 +Indole/26 −Indole; pCASPR: N = 2, 90 +Indole/26 −Indole) and motA (No-SCAR: N = 2, 16 positive/ 20 negative; pCASPR: N = 2, 16 positive/ 20 negative). Statistical significance was assessed using two-sided Fisher’s exact tests comparing editing efficiencies between No-SCAR and pCASPR methods for each target, with Holm–Bonferroni correction for multiple comparisons. **C)** Knock-in experiments to replace the *motAB* locus with mVenus in *Ec* RP437. Colony PCR results of screening 9 colonies from each plate and WT control excerpt from SI Fig. 3. The presence of a 2.8 kb band (top bands in gel) is indicative of WT *motAB* locus. The lower size band indicated by the yellow arrow (∼1.7 kb) indicates *mVenus* knock-in at the *motAB* locus and concomitant loss of the *motAB* genes. **D)** Quantification and comparison of fraction of positive edits obtained across all experimental replicates using no-SCAR (red) or pCASPR (grey) in *E. coli* RP437 as target strain for knockout (KO) and knock-in (KI) experiments at the *motAB* locus. The homology arm length for the KI construct was 140 bp per homology arm, compared to the 500 bp-long homology arms employed in *Ec* BL21 (panels A and B above) and the KO experiments. N = 5. Statistical testing was carried out using Fisher’s exact test: pCASPR significantly increased editing efficiency compared with the no-SCAR system for both motAB knockout (33.8% vs 11.9%, Fisher’s exact test, Holm-adjusted P = 0.0094) and mVenus knock-in (11.5% vs 0%, Fisher’s exact test, Holm-adjusted P = 0.0084).

A GFP-dropout pCASPR derivative (pCASPR-GFP:DO) was generated for guide-retargeting and visual monitoring of plasmid maintenance/curing (SI Fig. 4). In this derivative, the sgRNA N20 region is flanked by BsmBI Type IIS cloning sites so that target-specific spacer sequences can be exchanged locally without rebuilding the whole plasmid. Candidate plasmids were screened by loss of GFP signal after guide insertion and verified by sequencing across the guide region. Final plasmids were verified by whole-plasmid sequencing (Plasmidsaurus).

To make the Cas9-containing backbone compatible with Type IIS restriction enzymes, internal BsmBI and BbsI recognition sites in the Cas9 coding sequence were removed by synonymous substitution using synthetic DNA fragments and PCR assembly. PCR products were generated using high-fidelity DNA polymerase (Q5, NEB), separated by agarose gel electrophoresis, gel-purified, and assembled using BbsI-HF, BsmBI-v2 or BsmBI-HF, and T4 DNA ligase where applicable.

### pCASPR*-So* construction for *Shewanella oneidensis*

pCASPR*-So* was constructed by adapting the pCASPR single-plasmid architecture for *S. oneidensis* as described in SI Fig. 8. The spectinomycin-resistance marker was replaced with a kanamycin-resistance marker to generate an intermediate construct. The plasmid origin of replication region was then replaced with the Shewanella-compatible ColE1/OriTRP4 module. Finally, the guide N20 sequence was retargeted from the *E. coli motA* control sequence to a sequence targeting the *S. oneidensis mtrC* locus. The resulting pCASPR-*So*-*mtrC* plasmid retained the *tetR*/aTc inducible Cas9/sgRNA counterselection module, *araC*/pBAD-regulated lambda-Red genes, kanamycin selection, and the host-adapted replication/transfer module.

### Guide RNA design and cloning

Guide RNAs for pCASPR *E. coli* assays were designed against the *E. coli* BL21(DE3) genome using CHOPCHOP {Labun, 2019 #18} or Benchling (Biology Software, 2026), then screened for off-target using Cas-OFFinder {Bae, 2014 #19}. For each target, a 20 nt spacer adjacent to an NGG PAM was selected, and the PAM was omitted from the cloned guide sequence. The main pCASPR experiments used guide targets associated with *rpsL, lacZ/lacZY, trpD/trpLEDC*, and *motA*, together with a non-targeting control guide (pCASPR-GFP:DO).

For pCASPR Golden Gate guide cloning, complementary oligonucleotides were ordered with BsmBI-compatible overhangs for insertion into the GFP:DO cassette. Oligonucleotides were annealed at a final concentration of 1 μM in 1x T4 DNA ligase buffer by heating to 95°C followed by gradual cooling to room temperature. Golden Gate reactions contained 50 nM annealed oligonucleotide duplex, 50 ng pCASPR-GFP:DO, 5 units BsmBI-v2, and 200 units T4 DNA ligase in a 10 μL reaction. Reactions were cycled 20 times between 42°C and 16°C for 5 min at each temperature, followed by heat inactivation at 80°C for 20 min. Candidate guide plasmids were identified by loss of GFP signal and verified by Sanger sequencing across the guide region.

### Donor DNA construction

Linear dsDNA donor templates for pCASPR *motAB* editing were generated by PCR assembly of homology-arm fragments amplified from the target strain genome. For the *motAB* deletion donor, left and right homology arms were spliced to produce an approximately 1 kb donor that removed the *motAB* coding region without leaving an antibiotic-resistance marker. For fluorescent-protein knock-in donors, the homology arms were assembled with mVenus reporter sequence to replace the *motAB* locus; the mVenus donor shown in SI Fig. 6 is approximately 1.7 kb, and shorter variants (∼1 kb) were also produced using nested PCR primers. Donor assemblies were gel-purified and verified by Sanger sequencing across assembly junctions before editing.

For the multi-locus assays in *E. coli* BL21(DE3), donor DNA templates for the knock-outs (Δ*lacZY*, Δ*trpLEDC*, and Δ*motA*), which delete the entire coding sequence, were ordered as gBlocks (Integrated DNA Technologies, IDT) comprising 200-500 bp homology arms flanking the target deletion. gBlocks were amplified using high-fidelity polymerase, and gel-purified to generate sufficient quantity. The *rpsL* K43R donor was ordered as an 80 bp-long single-stranded oligonucleotide with four phosphorothioate bonds at the 5’ end (IDT), as described previously {Reisch, 2017 #6}(Supplementary Table 1).

For pCASPR-*So*-*mtrC* editing in *S. oneidensis*, the intended donor DNA introduced a SpyTag sequence adjacent to native *mtrC*. The donor design contained the SpyTag sequence flanked by mtrC/mtrA-adjacent homology regions and silent changes in the guide/PAM region to prevent Cas9 cleavage of the edited allele. A linear HDR donor strategy was tested first under the conditions described below. Because that strategy did not recover the desired edit, the repair template was subsequently assembled into pCASPR-*So*-*mtrC*-HDR, thereby coupling the donor template to the editing plasmid.

### pCASPR and no-SCAR functional assays and editing efficiencies

pCASPR-*motA* maintenance and induction were tested on non-selective LB, selective LB + L-arabinose and selective LB + L-arabinose and aTc (Fig. 1C). Growth under selection confirmed plasmid maintenance; loss of recovery after L-arabinose + aTc induction indicated donor-free Cas9/sgRNA counterselection at the chromosomal *motA* target. For editing, RP437 or BL21(DE3) cells harbouring either the pCASPR or no-SCAR plasmids were grown at 30°C to OD_600_ ≈ 0.4–0.5, induced with L-arabinose for 20 min to induce lambda-Red expression, made electrocompetent in ice-cold 10% glycerol, and transformed by electroporation (0.2 cm, 2.5 kV, ∼5 ms) with 200 ng linear donor DNA (unless otherwise stated) or 2 µM phosphorylated donor ssDNA. After recovery (2 h in LB at 30°C, or 5 h for *rpsL* K43R), cells were plated on selective aTc medium and screened by colony PCR or functional assays.

Editing efficiency was determined by screening surviving colonies using the method appropriate for each target. Edits at *motA* were confirmed by colony PCR. The *rpsL* K43R edit was assessed by growth in the presence and absence of streptomycin (50 μg/mL), and *lacZY* editing by blue-white screening on X-gal (40 μg/mL) and IPTG (0.1 mM). *trpLEDC* editing was assessed by growth on M9 medium in the presence and absence of 50 μM indole (*trpLEDC* deletion removes the ability to synthesise tryptophan, but edited cells retain the ability to convert indole to tryptophan, so edited clones grow on M9 with indole (but not without it). For the streptomycin, X-gal and indole screens, plates containing 50-100 colonies were assessed per transformation plate. Editing efficiency was calculated for each transformation and averaged across three independent transformations.

### Counterselection survival assays

Guide-dependent counterselection with pCASPR was tested in *E. coli* BL21(DE3) carrying a non-targeting (NT), *rpsL, lacZ*, and *motA* guide. Overnight cultures were grown at 30°C in selective LB, diluted to an OD_600_ of 0.1, then further diluted to 10^-3^, and 20 µL was spotted onto selective LB agar with the presence and absence of aTc (0, 50, 100, 200, 400 ng/mL).

BL21(DE3) cells harbouring either pCASPR or the no-SCAR plasmids were grown to saturation at 30°C in selective LB medium. Cultures were serially diluted ten-fold to 10^-11^ and 5 µL was spotted onto LB agar plates containing the appropriate antibiotics, with and without aTc. Colonies were counted at the lowest dilution at which individual colonies could be distinguished. Survival frequency was calculated as CFU/mL on +aTc divided by CFU/mL on -aTc plates, for three independent cultures.

### Plasmid curing

Temperature-dependent maintenance of pCASPR was tested using *E. coli* BL21(DE3) harbouring the pCASPR-GFP:DO plasmid. Cultures were grown to an OD_600_ of 0.1 at 30°C in selective LB, diluted 10^-3^ and 20 µL was spotted onto selective LB agar. Plates were incubated at 30, 37 or 42°C for 16-20 h.

To obtain strains cured of pCASPR, a single colony was inoculated into 3 mL of non-selective LB and grown at 42°C for 8 h (colony picked in the morning) or 16 h (colony picked in the afternoon). The culture was then streaked onto non-selective LB agar and incubated at 42°C for ∼16 h to obtain single colonies. Single colonies were then patched onto non-selective and selective LB agar, and the plates were incubated at 30°C for 20 h. Colonies that grew on non-selective LB but not on selective were scored as cured.

Due to the poor growth of Shewanella oneidensis at temperatures above 37°C, pCASPR-*So* could not be cured in a temperature-dependent manner as for *E. coli*.

### Data handling and statistics

Counterselection survival (CFU +aTc / CFU -aTc) and editing efficiency (edited/total screened) were analysed separately. Unedited survivors were counted only when genotyped as retaining the target allele, and “escaper” restricted to those cases or to no-donor survival experiments. Biological replicates were independent experiments from separate overnight cultures carried independently through induction, competent-cell prep, transformation, recovery, plating, and screening; technical plates aided countability only. For the counterselection survival assay statistical comparisons between pCASPR and No-SCAR were performed using unpaired Welch’s t-tests on log[[-transformed escape frequencies, with Holm correction for multiple comparisons. For the editing efficiency assay statistical significance was assessed using two-sided Fisher’s exact tests comparing editing efficiencies between No-SCAR and pCASPR methods for each target, with Holm–Bonferroni correction for multiple comparisons.

## Results

### Construction and functional validation of pCASPR

We first assembled pCASPR as a single-plasmid implementation of CRISPR-Cas9 counterselection and lambda-Red recombineering. The editing workflow retains the familiar CRISPR-recombineering logic: a target-specific sgRNA directs Cas9 to the chromosomal allele, unrepaired cleavage counterselects against the unedited chromosome, donor-mediated recombination repairs the target as either a scarless knockout or knock-in, and the editing plasmid is removed by temperature-sensitive curing (Fig. 1A; SI Fig. 1). The plasmid construct was built from no-SCAR-derived pCas9cr4 and pKDsgRNA modules following our earlier work editing the *motAB* locus of *E. coli* {Ridone, 2022 #20} to include the Cas9 cassette, sgRNA expression cassette, arabinose-inducible recombineering genes, antibiotic selection, and temperature-sensitive replication module on one plasmid (Fig. 1B). In the pCASPR-*motA* prototype, the pCas9cr4-derived fragment supplied the *tetR*/tetO-regulated *Cas9* module, while the pKDsgRNA-derived fragment supplied the sgRNA scaffold, *araC*/pBAD-inducible *gam-beta-exo* cassette, spectinomycin selection, and pSC101ts-type origin of replication. Agarose-gel analysis confirmed assembly fragments of the expected sizes and an approximately 12 kb NotI-linearised pCASPR product (SI Fig. 2A), and Sanger sequencing across the junction/guide region confirmed the intended assembly (SI Fig. 2B).

The assembled plasmid was then tested for the basic functions required for editing. Cells carrying pCASPR-*motA* grew on non-selective medium and under spectinomycin selection, including after arabinose induction of the lambda-Red cassette (Fig. 1C). In contrast, combined arabinose and aTc induction eliminated visible recovery under plasmid-selective conditions, consistent with induction of target-matched Cas9/sgRNA cleavage in the absence of a repair template (Fig. 1B). Cas9 activity was guide-dependent. A non-targeting guide retained growth after aTc induction, whereas guides targeting *rpsL, lacZ*, or *motA* reduced colony recovery after induction, showing that the counterselection phenotype required a chromosomal target for the expressed sgRNA (Fig. 1D). Additional aTc titration experiments showed that induction strength affected recovery and that the extent of growth inhibition differed between target guides, consistent with guide- and locus-dependent counterselection rather than a uniform toxicity response to aTc or Cas9 expression (SI Fig. 3).

**Figure 3.**
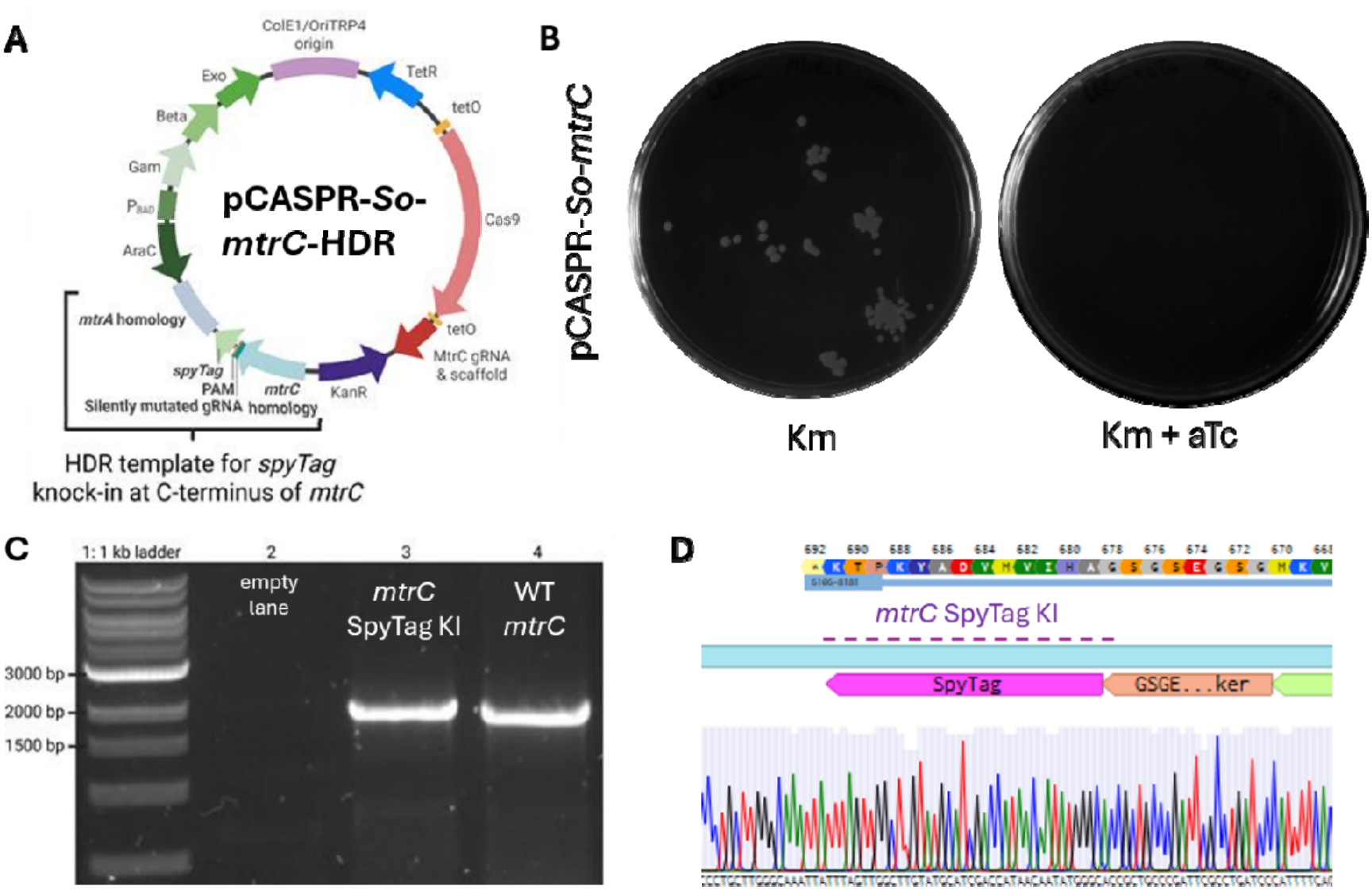
pCASPR*-So-mtrC-*HDR adapts the single-plasmid CRISPR/lambda-Red architecture for *Shewanella oneidensis* and enables SpyTag insertion at the native *mtrC* locus. **(A)** The pCASPR-*motA* backbone was modified by replacing spectinomycin resistance with kanamycin resistance, replacing the original replication/compatibility module with the ColE1/OriTRP4 module shown in the figure, and retargeting the sgRNA from the *E. coli motA* control sequence to the *S. oneidensis mtrC* locus. The donor DNA template for SpyTag Knock-in was also incorporated into the plasmid between the lambda-red and Kan^+^ resistance marker gene. The resulting *pCASPR-So*-*mtrC-*HDR plasmid retains the *tetR*/tetO-controlled Cas9 module, the *araC*/pBAD-controlled lambda-Red genes, kanamycin selection, and an *mtrC*-targeting sgRNA scaffold. The full construction path from pCASPR-*motA* to the *S. oneidensis* editor pCASPR*-So-mtrC-*HDR is shown in SI Fig. 8. **(B)** Testing counterselection in *S. oneidensis*. Photographs of LB Agar plates containing kanamycin (Km) +/-anhydrotetracycline (aTc), inoculated with *S. oneidensis* carrying pCASPR*-So-mtrC* (the pCASPR construct without the SpyTag donor DNA template incorporated). Targeting of the gRNA control sequence to *mtrC* results in cell death when induced with anhydrotetracycline, in absence of an HDR template. **(C)** PCR validation of candidate *mtrC*-SpyTag edited colonies. A colony PCR using genomic primers outside the donor region distinguished the edited allele from the unedited allele, with expected products of 1993 bp (lane 3) and 1930 bp (lane 4), respectively. **(D)** Sanger sequencing of colony-PCR amplicons confirmed SpyTag (purple) in-frame integration at the 5’ end of the intended *mtrC* locus (encoded on the lagging DNA strand) corresponding to the MtrC C-terminus.

To monitor plasmid curing and facilitate guide RNA exchange, we generated a GFP-marked, Golden Gate-compatible pCASPR derivative (pCASPR-GFP:DO). The GFP cassette, flanked by BsmBI sites upstream of the sgRNA scaffold, is replaced upon insertion of a new N20 spacer, providing a visual screen for successful guide cloning (SI Fig. S4). Hence, the pCASPR-GFP:DO plasmid was used as the non-targeting (NT) control plasmid in all counterselection assays. The temperature-sensitive maintenance module also behaved as expected. Clones carrying pCASPR-GFP:DO retained growth and fluorescence at 30°C and 37°C , but was not maintained at 42°C under selective conditions (Fig. 1E). Together, these assays established that the pCASPR backbone expressed all desirable features of the parental no-SCAR system (the guide-dependent counterselection, antibiotic selection, and temperature-sensitive plasmid curing) in a single construct.

### Comparison of editing performance between the single-plasmid and dual plasmid systems

We first evaluated the capacity of the single plasmid system (pCASPR) for donor-free counterselection across multiple genes and compared it to no-SCAR. For both systems, sgRNAs were directed against chromosomal targets including *rpsL, lacZ, trpD*, and *motA*. The non-targeting guide retained colony recovery across the aTc induction series, whereas target-matched guides reduced recovery after induction (Fig. 2A). Survival frequency varied between guides, with the no-SCAR *lacZ* guide showing the highest survival on aTc, whereas all pCASPR target-matched guides showed very low recovery across the guide set (Fig. 2A). These results demonstrate that pCASPR can be retargeted to several chromosomal loci and that the effectiveness of counterselection is guide-dependent. Because these assays measure colonies recovered after donor-free Cas9 induction, the conservative interpretation is counterselection survival rather than a mechanistic classification of every recovered colony as an escape mutant.

We tested whether retargeted pCASPR plasmids could produce edited clones (knockouts) after donor DNA transformation and compared the editing efficiency to no-SCAR. Linear dsDNA donors were used to generate Δ*lacZY* (, Δ*trpLEDC* and Δ*motA*, and single-stranded oligonucleotide donor was used to generate the *rpsL* K43R allele (Supplementary Table 1). Edited colonies were recovered at all four loci tested following screening by colony PCR (Fig. 2B, SI Fig. 5). The edited fraction among screened colonies was system- and locus-dependent. pCASPR gave a higher edited fraction than no-SCAR for *rpsL* K43R (p < 0.001), a similar fraction for Δ*motA* and a slightly lower fraction for Δ*lacZY* (p < 0.001; Fig. 2B). The two methods did not significantly differ in efficiency in deleting the *trp* operon (Δ*trpLEDC*). Thus, pCASPR supports editing at multiple *loci*, but its performance is not uniform across targets.

We also tested pCASPR and no-SCAR editing at the *motAB* locus in *E. coli* RP437 with two edits: a knockout (KO), in which *motAB* (1811 bp) was deleted, and a knock-in (KI), in which *motAB* was replaced with mVenus (1811 bp replaced by 717 bp; Fig. 2C, 2D). The mVenus donor was assembled into a 1.7 kb fragment from amplicons carrying flanking homology to *motAB* and the *mVenus* sequence and confirmed by Sanger sequencing (SI Fig. 6B). Nested PCR was used to generate a 1 kb-long version of the donor DNA from the assembled 1.7 kb fragment (SI Fig. 6C) of similar size to the KO donor DNA (1 kb). The knock-in donor DNA used shorter homology regions (140 bp) flanking the *mVenus* sequence compared to the knockout donor (500 bp; SI Fig. 6D). Candidate clones carrying a genomic replacement were identified by colony PCR and agarose gel electrophoresis (Fig. 2C, SI Fig. 7) and loss of motility by functional phenotyping using a soft-agar motility assay (SI Fig. 7D). The size of the colPCR amplicons was used to diagnose the genotype of each colony: WT *motAB* (2.8 kb), *motAB* KO (1 kb), *mVenus* KI at *motAB* locus (1.7 kb). The pCASPR system significantly increased editing efficiency compared with the no-SCAR system for both motAB knockout (33.8% vs 11.9%, P = 0.0094) and mVenus knock-in (11.5% vs 0%, P = 0.0084).

For the knockout, the proportion of positive clones among randomly selected colonies was similar for pCASPR and no-SCAR (Fig. 2D). For the knock-in, positive clones were recovered only with pCASPR, whereas none were detected with no-SCAR (Fig. 2C, 2D, SI Fig. 7). With pCASPR, knock-ins were recovered less frequently than knockouts. The pCASPR-GFP:DO plasmid was used as the non-targeting (NT) control plasmid in all counterselection assays.

### pCASPR-*So* adapts the single-plasmid architecture for *Shewanella oneidensis*

Finally, we asked whether the same editing logic could be rebuilt for a non-*E. coli* host. The *S. oneidensis* derivatives, (pCASPR*-So*) were built from the pCASPR-*motA* by replacing the spectinomycin resistance with kanamycin resistance, replacing the replication module with a ColE1/OriTRP4 module, and retargeting the sgRNA from the *E. coli motA* sequence to the native *S. oneidensis mtrC* locus (Fig. 3A, Fig. S8). The resulting plasmids, pCASPR-*So*-*mtrC*, retained the Cas9/sgRNA counterselection module and lambda-Red genes (Fig. 3A, SI Fig. 8B).

To test guide-dependent counterselection, we compared pCASPR*-So*-*motA*, in which the *motA* guide serves as a non-targeting control in this host, with pCASPR*-So*-*mtrC* (Fig. 3B, SI Fig. 8B). Both plasmids supported recovery on kanamycin without aTc. After aTc induction, pCASPR*-So*-*motA* retained colony recovery, whereas pCASPR*-So*-*mtrC* showed no recovery (Fig. 3B; SI Fig. 8B). This established that the host-adapted vector can express functional Cas9/sgRNA counterselection in *S. oneidensis* and that lethality depends on a chromosomal target.

We next attempted a SpyTag knock-in at the C-terminus of MtrC. A linear donor DNA did not yield edited colonies under the conditions tested, so the donor was cloned into the plasmid to give pCASPR-*So*-*mtrC*-HDR (Fig. 3A, SI Fig. 8A). This ensured that positive *So* transformants would also carry a copy of the HDR template alongside the pCASPR editing machinery. The donor carried the SpyTag sequence and silent changes in the guide/PAM region to protect the edited allele from re-cleavage. Colony PCR identified 4 of 4 colonies carrying the SpyTag edit (SI Fig. 9). Sanger sequencing of the amplicons confirmed in-frame SpyTag integration at the intended locus (1993 bp edited versus 1930 bp unedited allele; Fig. 3C, 3D). The temperature-sensitive RepA replication module of pCASPR was replaced in pCASPR-*So*, and *S. oneidensis* grows poorly at 42°C, so the plasmid could not be cured by a temperature shift. Despite multiple attempts to cure the pCASPR plasmid from Shewanella oneidensis, including up to four serial passages and repeated dilutions in antibiotic-free media, as well as overnight incubation at 37°C, the plasmid persisted in cells, indicating that these approaches were insufficient to achieve plasmid curing.

## Discussion

Here we have developed pCASPR, a compact single-plasmid platform for scarless genome engineering in *E. coli* and *S. oneidensis*. This retains the familiar workflow but in a single vector, executing Cas9/sgRNA counterselection, lambda-Red recombineering, antibiotic selection and, in *E. coli*, temperature-sensitive curing. Thus pCASPR removes the separate guide plasmid and the second round of CRISPR-mediated curing required by no-SCAR. In *E. coli*, this architecture enables retargetable, donor-templated editing across multiple loci and the recovery of plasmid-free edited strains. pCASPR is designed for scarless replacement of endogenous alleles by homologous recombination, and it complements insertion focused technologies rather than competing with them. Systems such as mini-Tn7/pGRG25 and pUF/pAIO enable site-specific integration, and RNA-guided transposons such as CAST and INTEGRATE can insert kilobase-scale or multiplexed cargo, but they add DNA scars at defined sites and do not replace endogenous sequence {Zhang, 1998 #2, Egger, 2020 #21; Gelsinger, 2024 #22; Klompe, 2019 #23; McKenzie, 2006 #24; Vo, 2021 #25}.

Because pCASPR is derived from no-SCAR components, the most suitable direct benchmark is no-SCAR itself. Importantly, our no-SCAR controls behaved as previously with similar survival frequencies despite differing guide targets {Reisch, 2015 #5}, and with a similar editing efficiency of *rpsL* K43R using the same guide and donor DNA {Reisch, 2017 #6}. As with no-SCAR, and common for CRISPR-based editing, pCASPR editing and survival were target-dependent across loci such as *rpsL, lacZY, trpLEDC*, and *motA*. pCASPR also recovered the mVenus knock-in at *motAB*, which was not recovered with no-SCAR under the same conditions. Overall, pCASPR offers several practical advantages over dual-plasmid systems, such as a lower survival frequency when no donor DNA is supplied and improved editing efficiencies.

The no-SCAR protocol requires the two plasmids to be introduced sequentially but carrying both modules on a single plasmid removes this step {Reisch, 2015 #5}. Leaky expression did not prevent construction or maintenance of pCASPR and we observed no growth defects in the absence of aTc, which may reflect the low copy number of the temperature-sensitive SC101 or tight TetR regulation of *cas9*. Curing is often an overlooked but essential step for resetting a strain for future editing or plasmid-based phenotyping. In the no-SCAR procedure, the pCas9cr4 plasmid is cured by a second round of Cas9-directed cleavage, using a pKDsgRNA plasmid targeting Cas9 to its own origin of replication. In pCASPR, the same plasmid that performs counterselection and recombination carries a temperature-sensitive replication module, a standard route for removing recombineering plasmids {Datsenko, 2000 #3; Reisch, 2017 #6}. Together, the single-transformation and single-step curing account for the time saving shown in Table 1.

**Table 1.**
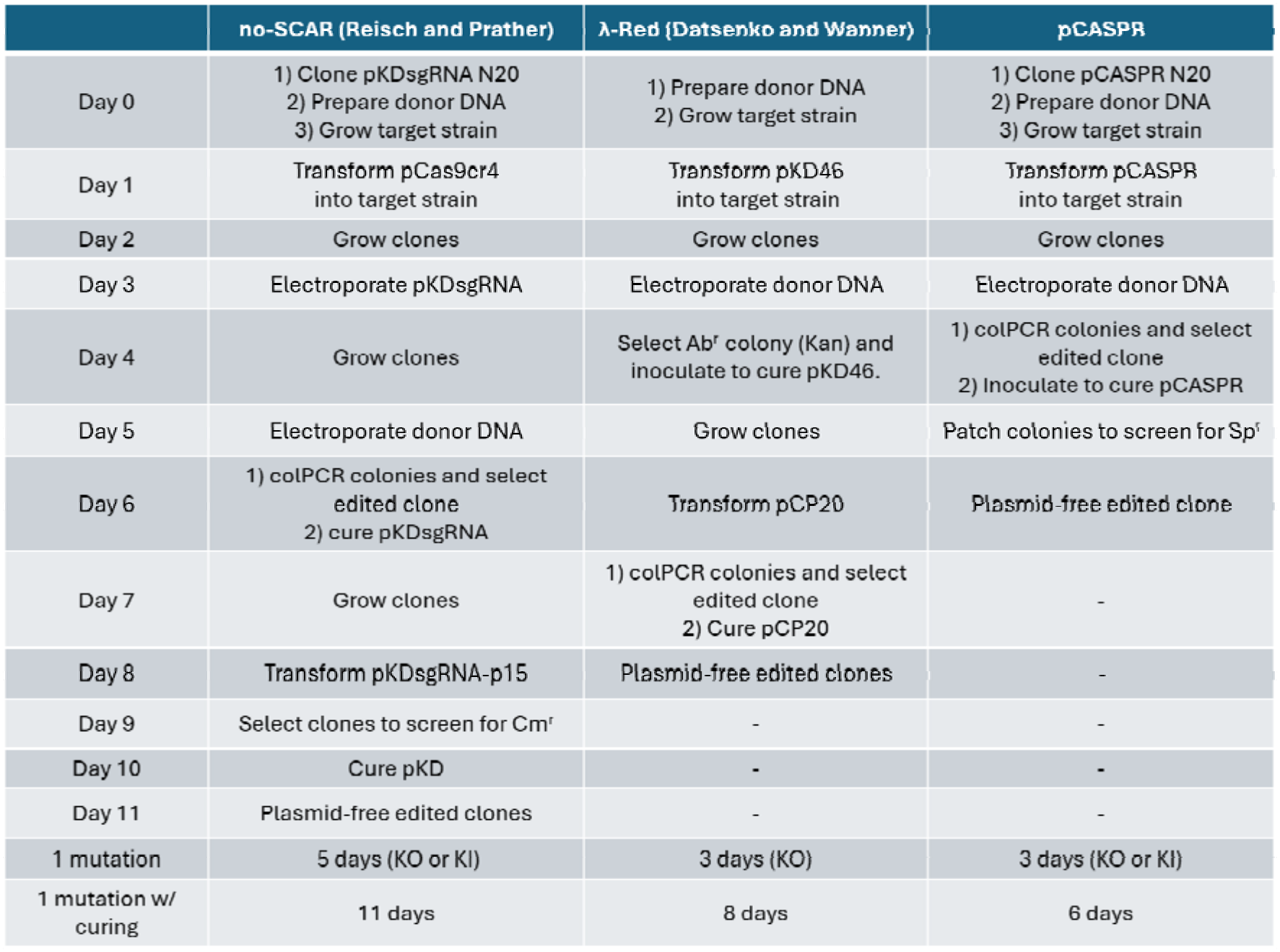
Comparison of gene editing methods for E. coli. Day-by-day breakdown of tasks involved in the no-SCAR method {Reisch, 2015 #5}, the Lambda-Red method {Datsenko, 2000 #3} and the pCASPR method presented here. The table assumes that all reagents are available on Day 0 with the goal to perform the first plasmid transformation on Day 1. Clone cultivation steps (‘Grow clones’) indicate overnight (∼16 hr) cultures at 30°C or 37°C depending on the plasmid used. Workdays represent 8 h shifts. This table was adapted from {Reisch, 2015 #5}.

Prior work in *S. oneidensis* showed that recombineering and CRISPR/Cas9 counterselection could be adapted to this organism, but those workflows depend on compatible replication, expression, selection, and transformation conditions {Corts, 2019 #14; Corts, 2019 #15}. pCASPR-*So*-*mtrC* shows that the single-plasmid architecture can be transferred to this host by exchanging the resistance marker and replication module. Use of a linear donor DNA fragment did not yield edited colonies, which we attribute to Cas9 cleavage and non-homologus end joining (NHEJ) prior to donor delivery. Cloning the donor into the plasmid, as others have done {Suzuki, 2020 #17}, resolved this but the extra cloning step runs against the ease-of-use goal that motivated pCASPR.

The temperature-sensitive RepA replication module was replaced in pCASPR-*So*, and *S. oneidensis* grows poorly at 42°C, preventing plasmid curation. Unlike previously reports {Suzuki, 2020 #17} serial passaging in antibiotic-free media did not result in the complete loss of plasmid. Future implementations could add autonomous or counterselectable curing modules, such as *sacB* for sucrose-sensitive lethality where it is host-compatible, so that plasmid loss occurs through active selection. Similarly, counterselection and editing efficiency in *E. coli* could be further improved by the use of transient RecA56-mediated suppression of the host SOS response {Moreb, 2017 #26}. This approach acts on the host response rather than on the editing architecture pCASPR.

In summary, pCASPR consolidates Cas9/sgRNA counterselection, lambda-Red recombination, guide exchange, selection and plasmid clearance into a single retargetable vector. In E. coli, it generated scarless edits across four tested loci with locus-dependent editing outcomes broadly comparable to no-SCAR, while simplifying transformation and curing. Rebuilding the replication and selection modules and incorporating the donor template enabled a proof-of-concept SpyTag insertion in S. oneidensis, but also showed that donor delivery and plasmid clearance remain host-specific design problems. pCASPR therefore provides a practical modular framework for single-plasmid bacterial genome editing whose performance and curing strategy should be validated in each new host.

