## Supplementary Figs S1-S9 for "pCASPR: A Single-Plasmid Platform for Scarless Bacterial Genome Editing"

### SUPPLEMENTARY FIGURES.

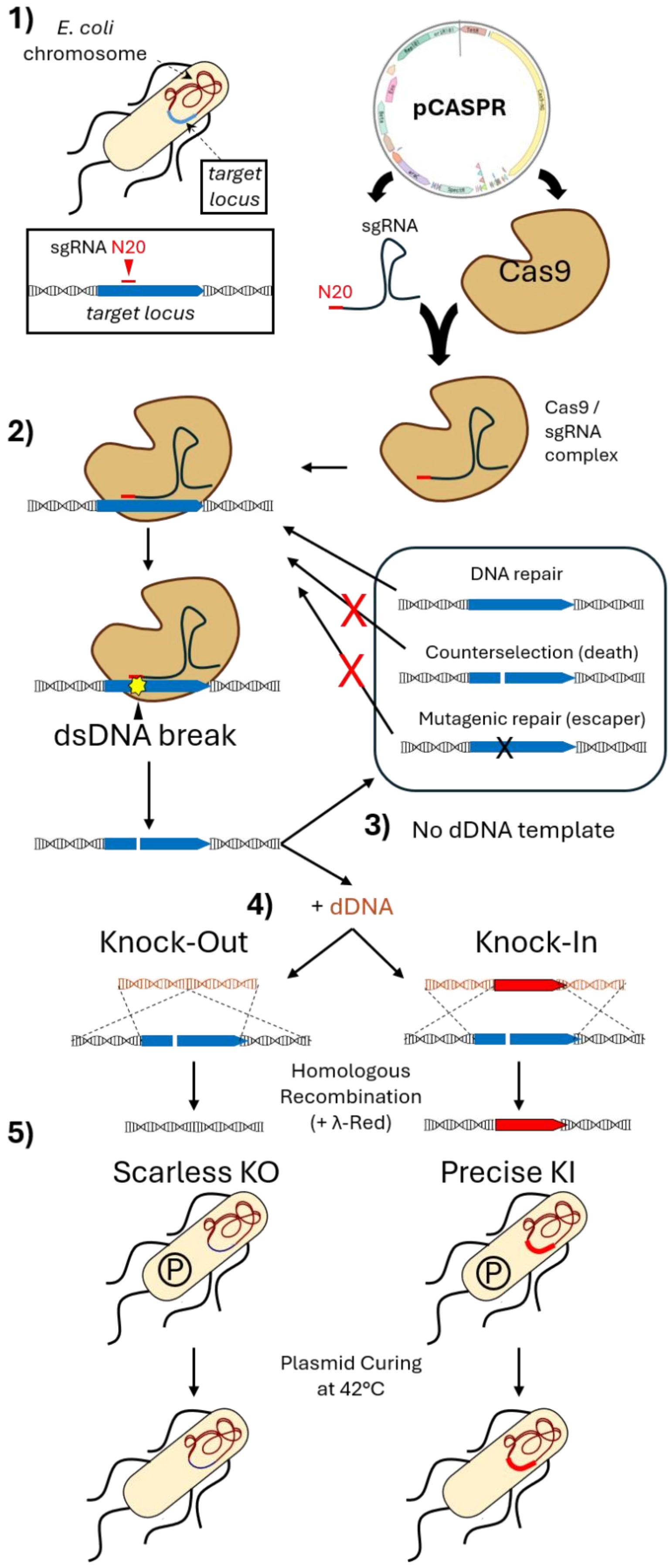

**Supplementary Figure 1. Mechanism of gene editing using a single plasmid system.**
1)The target cell is transformed with pCASPR to coexpress Cas9 and a single guide RNA able to hybridize to the target locus via its N20 sequence. 2) Activation of the Cas9/sgRNA complex causes a double stranded DNA break at the target locus. 3) In absence of a dsDNA repair template, the cell can repair the DNA break, regenerating the N20 site. This enables Cas9/sgRNA to cut the chromosome once more and the cycle repeats until cell death (counterselection) or N20 or PAM sites are mutated via error prone DNA repair (mutagenic repair), causing loss of function of the CRISPR system. This can lead to the propagation of false positive clones which survived counterselection. 4) In the presence of an appropriate dsDNA template the locus is repaired via Homologous Recombination. The process is enhanced by co-expression of the λ-RED machinery. The dsDNA template of choice determines the editing outcome (knockout or knock-in). The loss of N20 target upon recombination prevents further Cas9 activity. Edited cells are re-sequenced at their target locus to isolate the correct clone. 5) The editing plasmid can be removed from the edited cells by growing clones at 42°C. The temperature-sensitive origin of replication prevents plasmid propagation at high temperatures to regenerate a plasmid-free edited strain.

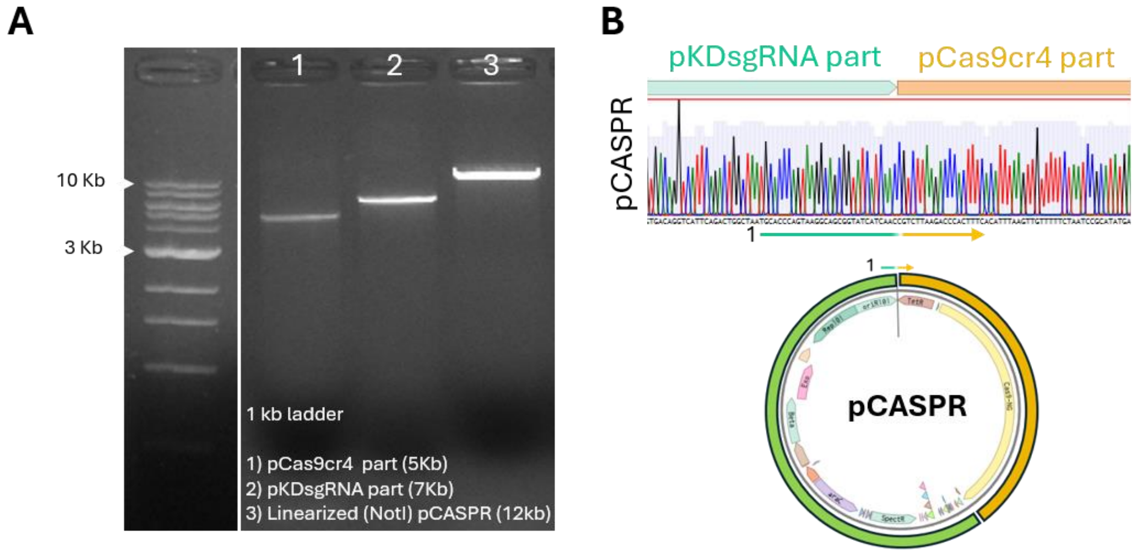


**Supplementary Figure 2. Construction of a single plasmid for scarless recombineering.**
(A) Agarose-gel validation of the assembly fragments and completed construct. Lane M, 1 kb DNA ladder; lane 1, pCas9cr4-derived fragment, approximately 5 kb; lane 2, pKDsgRNA-derived fragment, approximately 7 kb; lane 3, NotI-linearized pCASPR, approximately 12 kb. (B) Sequence validation of the assembled pCASPR junction/guide region, confirming the expected fusion between the pKDsgRNA-derived and pCas9cr4-derived modules.

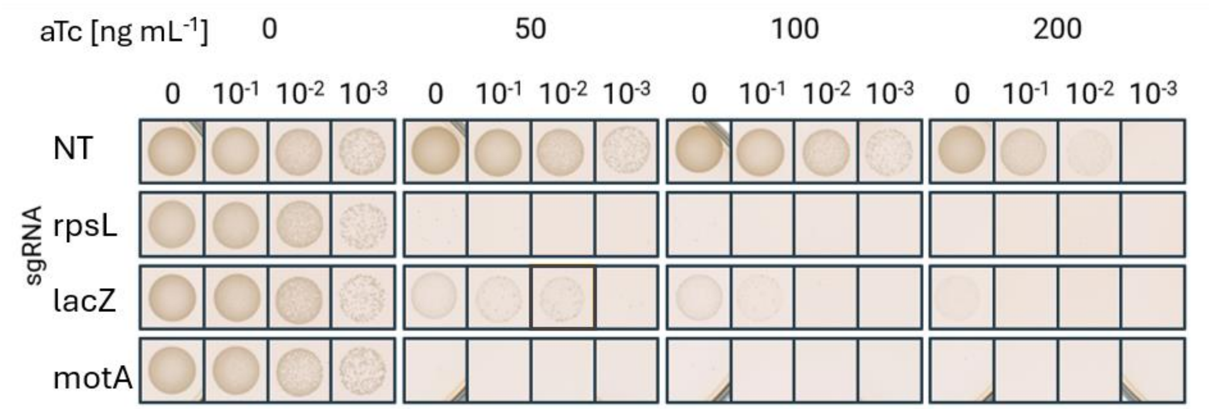


**Supplementary Figure 3. Testing and optimization of CRISPR activation using aTC.** Complete testing of aTc-inducible Cas9/sgRNA activity tested with a non-targeting (NT) guide (pCASPR-GFP:DO) and guides directed against *rpsL*, *lacZ*, or *motA* across increasing amounts of aTc (0, 50, 100 and 200 ng/mL). No colonies were observed when cells were cultured on 400 ng/ml aTc (data not shown). Dilution factors for inoculums are provided above each column.

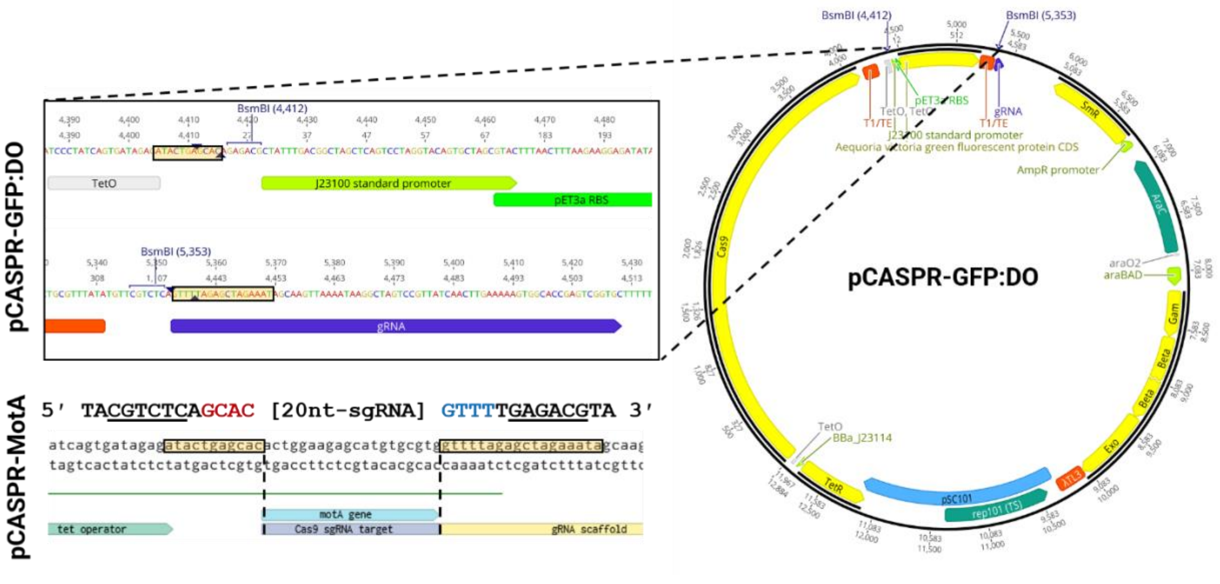


**Supplementary Figure 4. Map and sequences of interest of pCASPR-GFP:DO.** This GFP-drop out variant of the pCASPR system was designed for facilitated retargeting of the system and monitoring of the curing process. The pCASPR-GFP:DO carries a GFP expression cassette flanked by BsmBI cloning sites (highlighted by the orange boxes) in place of the N20 sgRNA-targeting sequence. Constitutive expression of GFP helps screen for correct clones when replacing the N20 sequence of pCASPR; loss of fluorescence (GFP drop-out) in a clone indicates successful integration of a new N20 sequence in pCASPR. The temperature-dependent curing of the plasmid can be monitored by loss of fluorescence. Complete loss of GFP emission indicates successful curing of the pCASPR plasmid.

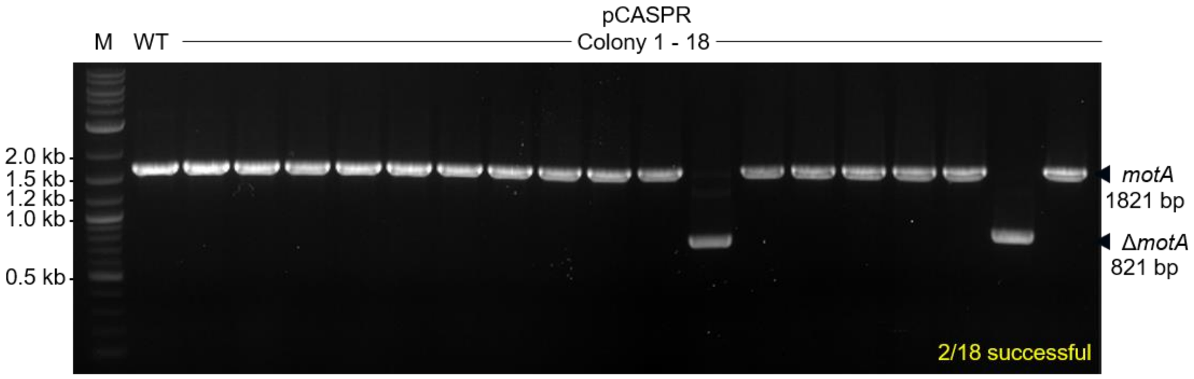


**Supplementary Figure 5. Example of colony PCR screening of pCASPR-mediated editing at the *motA* locus in BL21(DE3).** Colony PCR was performed using primers flanking the *motA* locus to distinguish edited from unedited colonies by amplicon size. 2-log DNA ladder (M) and wild-type control (WT) included per experiment. The editing efficiency of this knock-out experiment was 2/18.

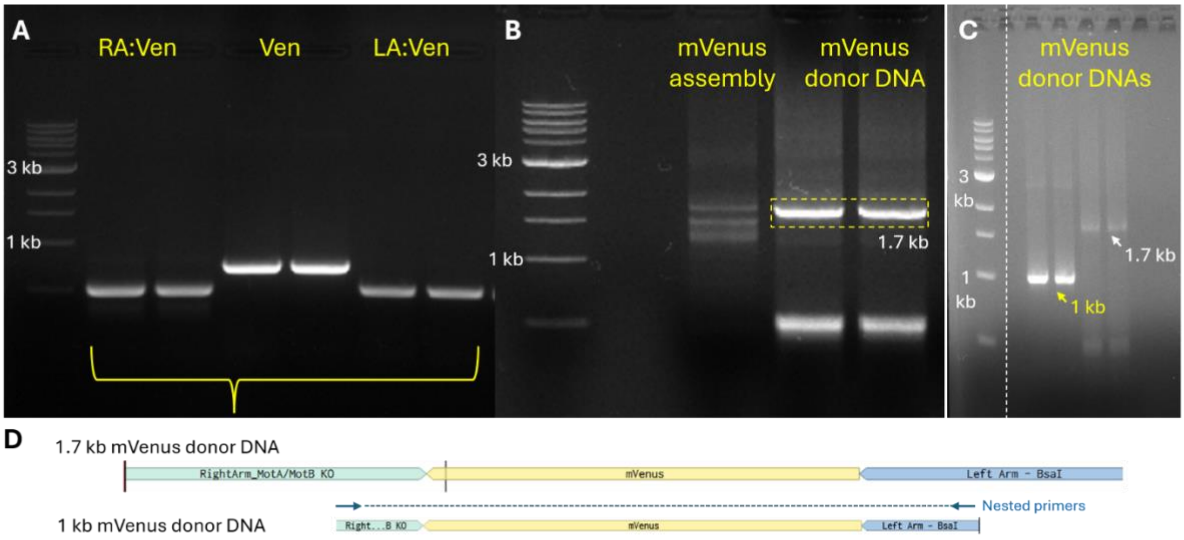


**Supplementary Figure 6. Assembly and use of donor DNA cassette to replace the genomic copy of the *motAB* genes with *mVenus*.** A) agarose gel of Right homology arm (RA:) and Left homology arms (LA:) fragments, their insert gene for mVenus (Ven) generated for assembly of donor DNAs. B) products resulting from the three-way assembly of donor DNA fragments and further amplification of the 1.7 kb donor cassette. C) Nested PCR product of the 1 kb mVenus donor DNA used in the editing experiments (yellow arrow), next to its 1.7 kb template (white arrow). The 3 kb and 1 kb markers are labelled on each gel. D) Schematic diagrams of the structure of the longer mVenus donor DNA assembly (1.7 kb) and its nested 1 kb amplicon that was used in the knock-in experiments.

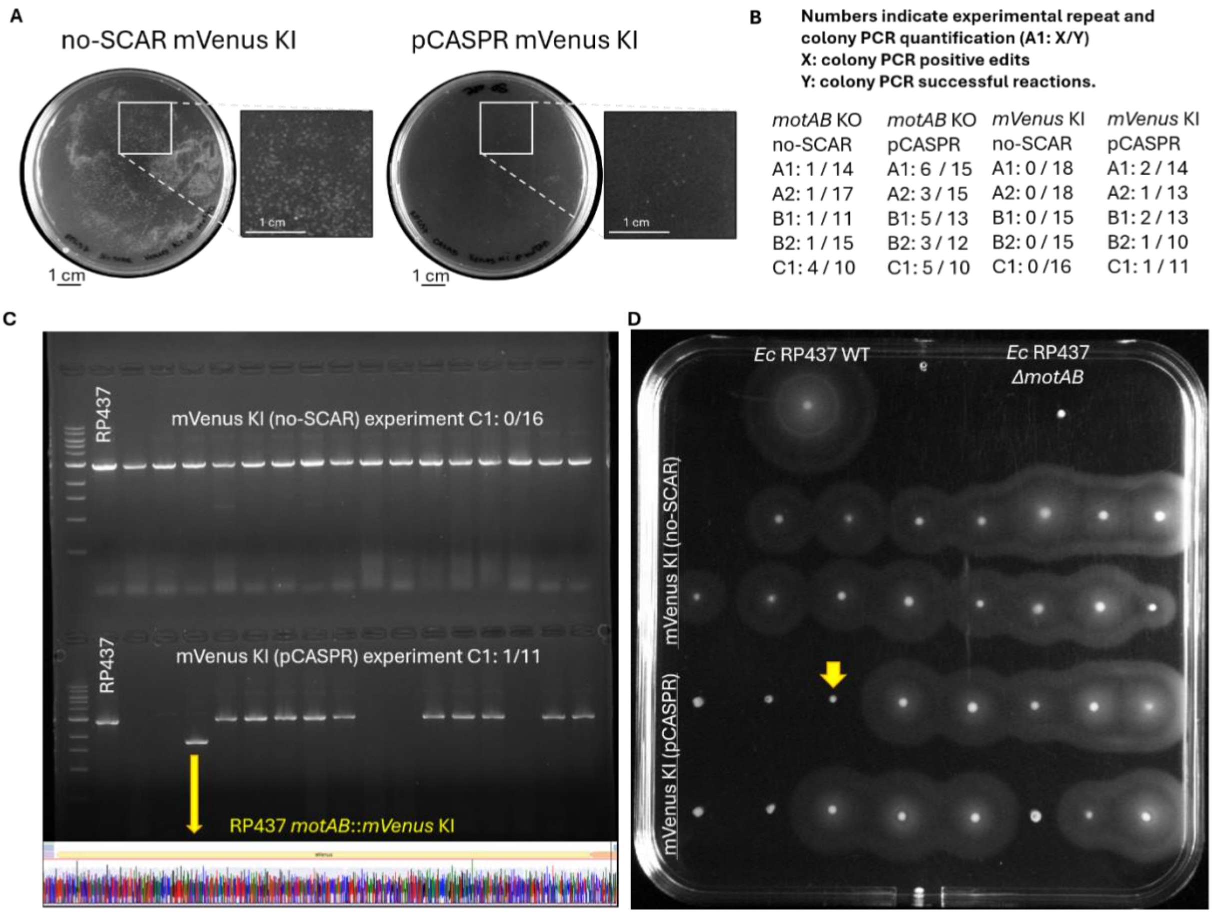


**Supplementary Figure 7. Comparing knock-in efficiency between no-SCAR and pCASPR systems to replace the genomic *motAB* locus of *Ec* RP437 with *mVenus*.** A) Counterselection plates containing aTc after electroporation of the mVenus dDNA in *Ec* RP437 carrying either the no-SCAR (left) or pCASPR (right) systems. Insets show density of candidate edited clones after plating the same volumes (50 µl). B) quantification of editing efficiency using colony PCR targeting the *motAB* locus across 5 replicate experiments (A1,A2, B1,B2,C1) of KO and KI using both systems. Numbers (x / y) indicate the colony PCR positive edits (x) detected amongst the successful colony PCR reactions (y) in the screening panel, plotted in Fig. 2D. C) Agarose gel elecrophoresis of Colony PCR screening panel for 1 control WT RP437 colony (~2.8 kb amplicon) and 16 candidate clones from mVenus KI using no-SCAR (top) or using pCASPR (bottom). Marker ladder is 1 kb from NEB. The yellow arrow points at Sanger sequencing results for the *mVenus*-positive PCR amplicon (~1.7 kb). The chromatogram below indicates successful in-frame replacement of the *motAB* locus with the *mVenus* gene. D) Screening edited cells for disruption of motility after gene editing at the *motAB* locus. Soft-agar motility assay was performed for each of the clones subjected to colony PCR in panel C. One colony out of the 16 from the no-SCAR experiment failed to grow (top left); all other colonies in the same group were motile and unaffected by the no-SCAR procedure. The yellow arrow indicates the colony edited as intended using the pCASPR system and confirmed via Sanger in panel C. While all unedited colonies showed swim rings, some colonies that failed the PCR assay also displayed disrupted motility, but were not characterized further.

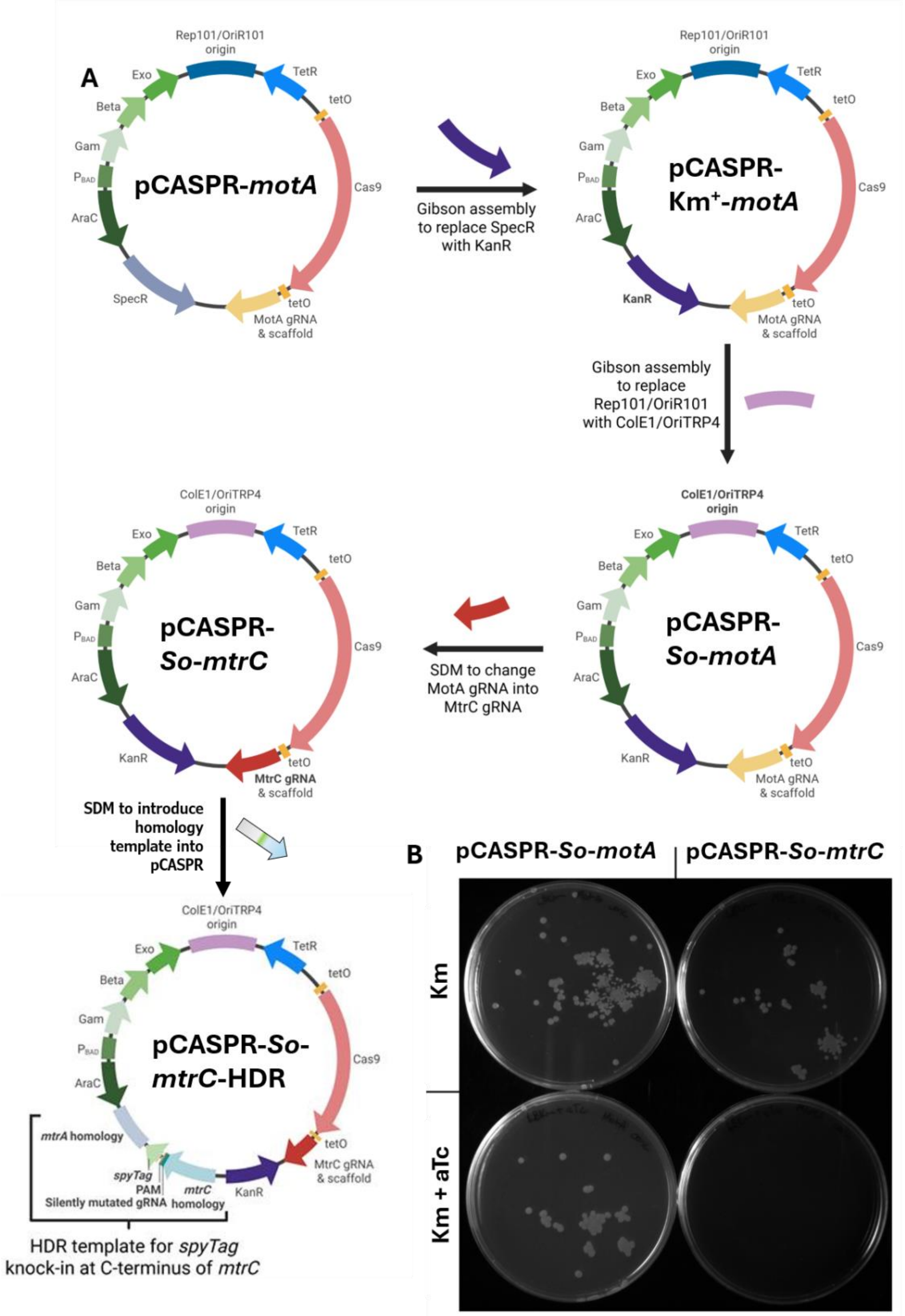

A
Rep101/OriR101 origin
Exo
TetR
tetO
Beta
Gam
P_BAD
pCASPR-motA
Cas9
AraC
SpecR
MotA gRNA & scaffold
Gibson assembly to replace SpecR with KanR
pCASPR-Km+-motA
KanR
Gibson assembly to replace Rep101/OriR101 with ColE1/OriTRP4
ColE1/OriTRP4 origin
pCASPR-So-motA
SDM to change MotA gRNA into MtrC gRNA
pCASPR-So-mtrC
MtrC gRNA & scaffold
SDM to introduce homology template into pCASPR
pCASPR-So-mtrC-HDR
mtrA homology
spyTag
PAM
Silently mutated gRNA
mtrC homology
HDR template for spyTag knock-in at C-terminus of mtrC
B
pCASPR-So-motA
pCASPR-So-mtrC
Km
Km + aTc

**Supplementary Figure 8. Development and validation of a pCASPR variant for use in *Shewanella oneidensis*.** A) First, the antibiotic resistance marker (SpecR) of pCASPR-*motA* was replaced with KanR to yield pCASPR-Km$^{+}$-*motA*. Then the origin of replication was replaced to facilitate maintenance in *Shewanella* (pCASPR-*So-motA*). Then, the N20 region of the sgRNA-coding section was replaced by site-directed mutagenesis (SDM) with an N20 targeting the *mtrC* gene in *S. oneidensis* (pCASPR-*So-mtrC*). This construct was used to validate counterselection and assess CRISPR activity at the *mtrC* locus. Finally, the donor DNA for genomic knock-in of the SpyTag adjacent to native *mtrC*, was incorporated into the pCASPR-*So-mtrC* plasmid. The donor design contains SpyTag sequence flanked by *mtrC* and *mtrA* homology regions and silent changes in the guide/PAM region to prevent re-cleavage by the Cas9/sgRNA complex after editing. This was installed between the λ-Red and Kan$^{+}$ genes and the construct was named pCASPR-*So-mtrC*-HDR. The linear HDR template strategy was unsuccessful under the conditions tested, consistent with genome cleavage occurring before donor delivery. **(B)** Target-dependent counterselection in *S. oneidensis*. Cells carrying pCASPR-*So-motA*, which serves as a non-targeting control in *S. oneidensis*, or pCASPR-*So-mtrC* were plated on kanamycin medium with or without aTc. Both constructs produced colonies under kanamycin selection without aTc. Upon aTc induction, colony recovery was retained for the *motA*-control guide but strongly reduced for the *mtrC*-targeting guide, indicating functional Cas9/sgRNA counterselection in *S. oneidensis*.

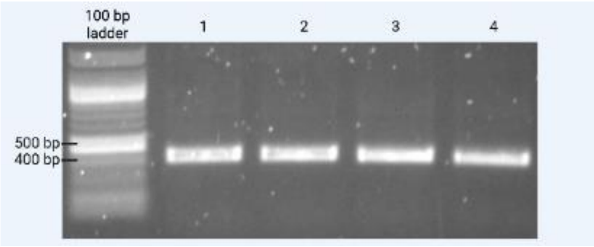


**Supplementary Figure 9. Colony PCR genotyping of candidate Shewanella clones after SpyTag Knock-In via pCASPR-*So-mtrC*-HDR.** Gel electrophoresis of colony PCR amplicons from four candidate edited Shewanella clones. The primer pair used annealed to the genome (forward primer) and the SpyTag (reverse primer). A 400 bp amplicon indicated the successful integration and correct orientation of the SpyTag coding sequence at the *mtrC* gene. Marker lanes corresponding to 400 bp and 500 bp are highlighted on the 100 bp ladder.

**Supplementary Tables.**

Table S1: Plasmids, N20 targeting sequences and donor DNA sequences
Table S2: Oligonucleotide primers
